# “3G” Trial: An RNA Editing Signature for Guiding Gastric Cancer Chemotherapy

**DOI:** 10.1101/2020.06.22.164038

**Authors:** Omer An, Yangyang Song, Xinyu Ke, Jimmy Bok-Yan So, Raghav Sundar, Henry Yang, Sun Young Rha, Lee Ming Hui, Tay Su Ting, Ong Xue Wen, Angie Tan Lay Keng, Matthew Chau Hsien Ng, Erwin Tantoso, Leilei Chen, Patrick Tan, Wei Peng Yong, Singapore Gastric Cancer Consortium (SGCC)

## Abstract

**Background & Aims:** Gastric cancer (GC) cases are often diagnosed at an advanced stage with poor prognosis. Platinum-based chemotherapy has been internationally accepted as first-line therapy for inoperable or metastatic GC. To achieve greater benefits, it is critical to select patients who are eligible for the treatment. Albeit gene expression profiling has been widely used as a genomic classifier to identify molecular subtypes of GC and stratify patients for different chemotherapy regimens, the prediction accuracy remains to be improved. More recently, adenosine-to-inosine (A-to-I) RNA editing has emerged as a new player contributing to GC development and progression, offering potential clinical utility for diagnosis and treatment.

**Methods:** We conducted a transcriptome-wide RNA editing analysis of a cohort of 104 patients with advanced GC and identified an RNA editing (GCRE) signature to guide GC chemotherapy, using a systematic computational approach followed by both *in vitro* validations and *in silico* validations in TCGA.

**Results:** We found that RNA editing events alone stand as a prognostic and predictive biomarker in advanced GC. We developed a GCRE score based on the GCRE signature consisting of 50 editing sites associated with 29 genes and achieved a high accuracy (84%) of predicting patient response to chemotherapy. Of note, patients demonstrating higher editing levels of this panel of sites present a better overall response. Consistently, GC cell lines with higher editing levels showed higher chemosensitivity. Applying the GCRE score on TCGA dataset confirmed that responders had significantly higher levels of editing in advanced GC.

**Conclusions:** Overall, the GCRE signature reliably stratifies patients with advanced GC and predicts response from chemotherapy.

**Significance:** Despite the increasing documentation of RNA editing and its functional regulation, the translational potential of RNA editome in cancer remains largely under-investigated. This study reports for the first time an RNA editing signature in advanced GC, to reliably stratify patients with advanced disease to predict response from chemotherapy independently of gene expression profiling and other genomic and epigenetic changes. For this purpose, a bioinformatics approach was used to develop a GCRE score based on a panel of 50 editing sites from 29 unique genes (GCRE signature), followed by an experimental evaluation of their clinical utility as predictive biomarker in GC cell lines and *in silico* validation in using RNA sequencing (RNA-Seq) datasets from TCGA. The applied methodology provides a robust means of an RNA editing signature to be investigated in patients with advanced GC. Overall, this study provides insights into the translation of RNA editing process into predictive clinical applications to direct chemotherapy against GC.

## Introduction

Gastric cancer is the third leading cause of cancer death worldwide responsible for more than 780,000 deaths annually^1, 2^. Asian population carries a higher risk for the disease in terms of incidence and mortality. The current treatment options for the advanced stage mainly employ palliative chemotherapy based on fluoropyrimidine and platinum-based compounds. However, an important question is whether we can improve the selection of patients with advanced GC for chemotherapy in order to achieve greater benefit from the treatment. To address this, different groups have put much effort in stratifying patients with GC into molecular subtypes^3–9^; however, these studies heavily focused on gene expression and/or mutation profiling, often with the limitations of microarray-based platforms or small sample sizes, thus novel molecular data types and approaches are needed for better guiding the chemotherapy treatment for this class of patients. Here, we demonstrate A-to-I RNA editing as a novel epigenetic classifier to reliably identify responders to chemotherapy.

RNA editing is a post- and/or co-transcriptional modification that results in specific nucleotide changes that occur on the RNA. In humans, the most frequent type of RNA editing is the conversion of adenosine to inosine (A-to-I), which is catalysed by ADAR proteins. In vertebrates, a family of 3 ADARs, ADAR1, ADAR2 and ADAR3, has been characterized^10^. ADAR1 and ADAR2 catalyse all currently known A-to-I editing sites. Inosine (I) essentially mimics guanosine (G), therefore ADAR proteins introduce a virtual A-to-G substitution in transcripts. Such changes can lead to specific amino acid substitutions^11–16^, alternative splicing^17^, microRNA-mediated gene silencing^18, 19^, or changes in transcript localization and stability^20–22^.

As reported by us and others in the past decade, dysregulated A-to-I editing is a key driver in the pathogenesis of various cancers, such as breast cancer^23^, glioma^24, 25^, chronic myeloid leukaemia^26^, hepatocellular carcinoma^11, 27^, and esophageal squamous cell carcinoma^12^. Our group provided the first extensive transcriptome-wide RNA editing analysis of primary gastric tumors and highlighted a major role for RNA editing in GC disease and progression^28^. This observation has been missed by previous next generation sequencing analyses of GC focused on DNA alterations alone. We reported that GC displays a severely disrupted RNA editing balance induced by the differentially expressed ADARs (ADAR1 and ADAR2). Clinically, the differentially expressed ADARs, which are characterized by ADAR1 overexpression and ADAR2 downregulation in tumors, have great prognostic value and diagnostic potential for primary GC. However, the role of RNA editing in inoperable, locally advanced or recurrent and/or metastatic GC and whether RNA editing signature can be used to prospectively and retrospectively stratify patients with advanced GC and predict response from chemotherapy, remain largely unknown.

Despite the fact that ADARs are responsible for A-to-I RNA editing activity, there is not always a linear relationship between the expression and activity of ADARs and editing frequencies of their target RNAs^29–31^, due to their differential subcellular distribution^32^, *cis*- and *trans*-regulatory interactors^33–36^ and post-transcriptional modifications^30, 37^. On the other hand, changes in the editing level of individual sites have been shown to play a driver role in several cancer types^11, 24, 38^. Therefore, RNA editing events are considered as a better proxy than ADAR expression *per se* to provide molecular information to be translated into clinical applications. We have previously initiated a translational “3G” trial to investigate the benefit of using a genomic classifier to guide the choice of two platinum-based chemotherapy regimens in an advanced GC setting^7^. In this study, we conducted a high-throughput RNA sequencing (RNA-Seq) analysis of 104 patients with advanced GC who had been enrolled into the “3G” trial and investigated the clinical utility of RNA editing events in advanced GC. To our knowledge, this is the first report which demonstrates that RNA editing alone can be employed as a prognostic and predictive factor in advanced GC, and more importantly, a panel of 50 editing sites could be readily detected in patients with advanced GC and accurately predicts outcome of chemotherapy. Overall, our study provides insight into the role of RNA editing in GC, which may facilitate the therapeutic decision making.

## Results

### The landscape of A-to-I RNA editing in advanced GC tumors

We conducted a genome-wide A-to-I RNA editing analysis using RNA-Seq data of endoscopic tumor biopsies obtained from 104 patients with advanced, metastatic or recurrent GC prior to their first-line palliative chemotherapy (platinum-fluoropyrimidine doublet chemotherapy regime) (**Supplementary Table S1**). Applying our established RNA editing pipeline^34, 39^ with stringent filtering criteria (**Materials and Methods**), we identified a total of 2,154,091 high confidence A-to-I RNA editing sites, with a median number of 17,000 editing sites per sample (**Figure 1A**), predominantly located in introns and 3 ’untranslated regions (**Figure 1B**), consistent with the previous reports^28, 40^. The number of editing sites moderately correlated with the total number of sequencing reads (Pearson’s r = 0.41, p = 1.96e-05) and with the overall editing activity (r = 0.39, p = 3.87e-05), where the latter was assessed by Alu Editing Index^41^. The overall editing activity, however, was independent of the total number of sequencing reads (r = −0.03, p = 0.8) and comparable across the samples (range 0.77-1.^34^, average = 1.03, stddev = 0.13, excluding 1 outlier). The distribution of the number of editing sites across the samples revealed an overwhelming number of sample-specific sites (n = 159,146), as well as 780 shared sites referred to as hotspots (i.e. edited in all the samples in the cohort) (**Figure 1C** and **Supplementary Table S2**). We included these hotspot editing sites for further analysis.

**Figure 1.**
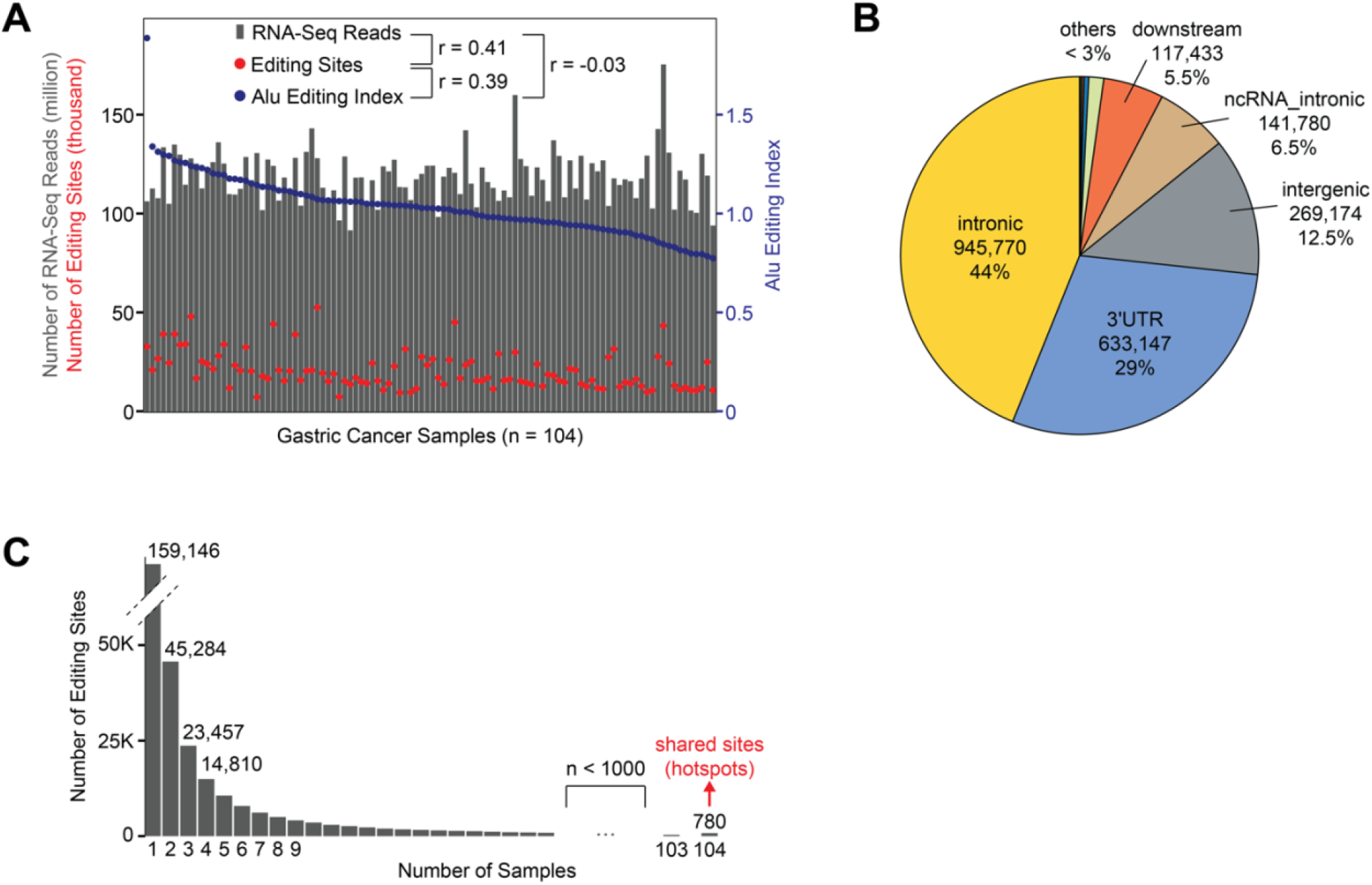
A-to-I RNA editing landscape in advanced GC cases. (*A*) Number of RNA sequencing reads, number of editing sites and Alu Editing Index (AEI) of 104 GC samples. The samples are sorted based on the decreasing value of AEI. r = Pearson correlation coefficient. (*B*) Distribution of 2,154,091 high confidence editing sites over annotated genomic regions. others = ncRNA_exonic, upstream, 5’UTR, exonic, upstream;downstream, splicing. (*C*) Distribution of number of editing sites across the samples, where shared sites that are edited in all the samples in this cohort (hotspots) are highlighted.

### RNA editing is a prognostic marker in advanced GC

First, we queried whether RNA editing has a prognostic value in advanced GC. To this end, we performed an unsupervised k-means clustering based on the RNA editing levels in an unbiased manner, i.e. including all the hotspot editing sites (n = 780) identified from the RNA-Seq data and all the patients with available survival data (n = 54). This resulted in two distinct clusters, which we defined as “high editing cluster” and “low editing cluster” referring to the relative average editing levels in the clusters (**Figure 2A**). The high editing cluster demonstrated significantly better patient survival (p = 0.037, k = 2) (**Figure 2B**), which was also evident when using hierarchical clustering (p = 0.067) (**Supplementary Figure 1**). Instead, the baseline patient characteristics we investigated did not correlate with the patient survival (Cox Proportional-Hazards univariate analysis, **Supplementary Table S3**).

**Figure 2.**
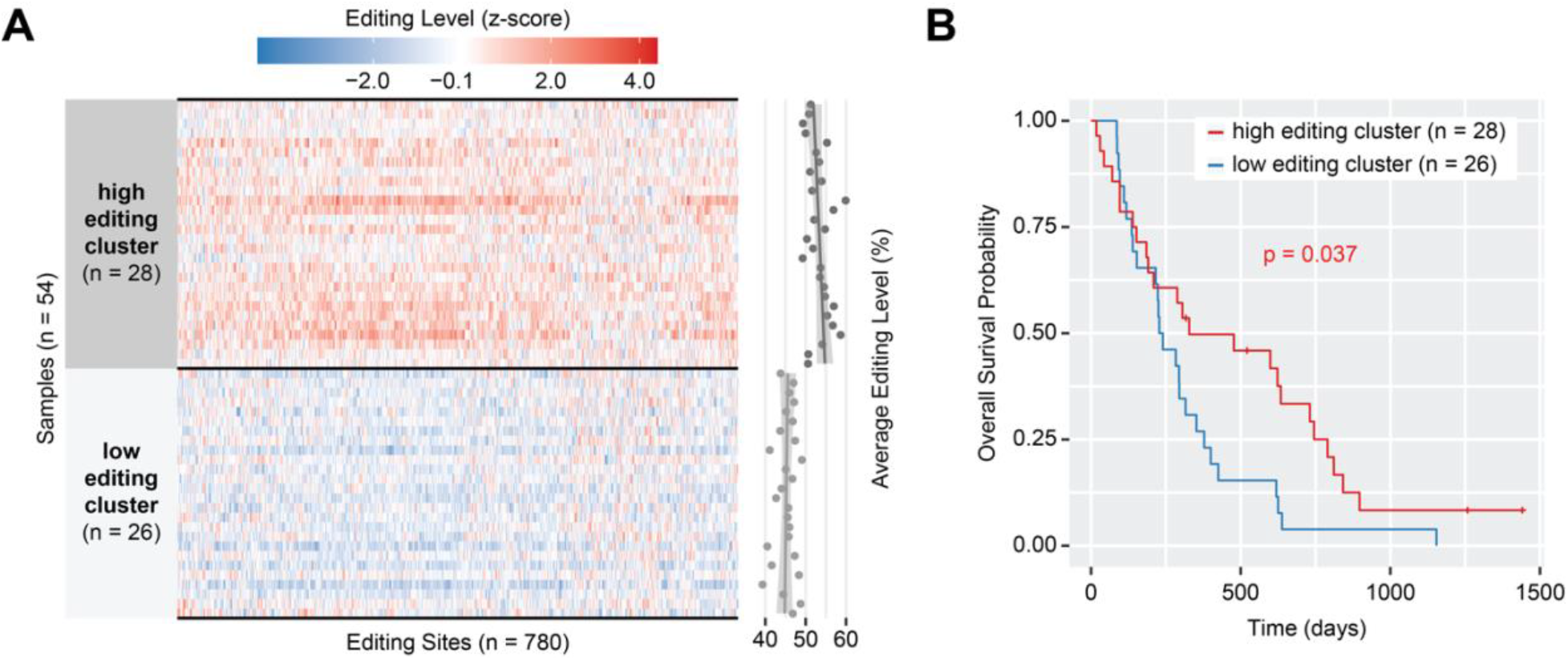
RNA editing hotspots as a prognostic marker in advanced GC. (*A*) Unsupervised k-means (k = 2) clustering of advanced GC samples based on RNA editing levels of 780 hotspot editing sites. The scatterplot shows average editing levels per sample. Of the cohort, 54 samples with survival data available are included in the analysis. (*B*) Survival plot of the two editing clusters. Data with hierarchical clustering is given in **Supplementary Figure 1**.

### RNA editing is a predictive marker in advanced GC

To investigate whether RNA editing also has a predictive value in GC, we focused on the overall response data of the patients to palliative platinum-based chemotherapy. For each of 780 hotspot editing sites, we applied Pearson correlation test between the RNA editing levels and overall response to chemotherapy (**Materials and Methods**). This led us to identify 53 key editing sites which showed significant correlation (p < 0.05) (**Figure 3A** and **Supplementary Table S4**). Interestingly, 50 of these sites had a positive correlation, implying the higher the editing level the better the response (response categories are numerically represented as PD = 0, SD = 1, PR = 2). Re-applying clustering on the patients by using the editing levels of only these 53 sites resulted in 75% accuracy of predicting the responders (i.e. patients who achieved PR) (**Figure 3B**), which was a significantly better prediction compared to a random selection of 53 sites (empirical p-value = 0.00409, N = 100,000, **Supplementary Figure 2**). Randomization test thus supports the validity of our selection method and highlights the importance of these sites as predictive markers.

**Figure 3.**
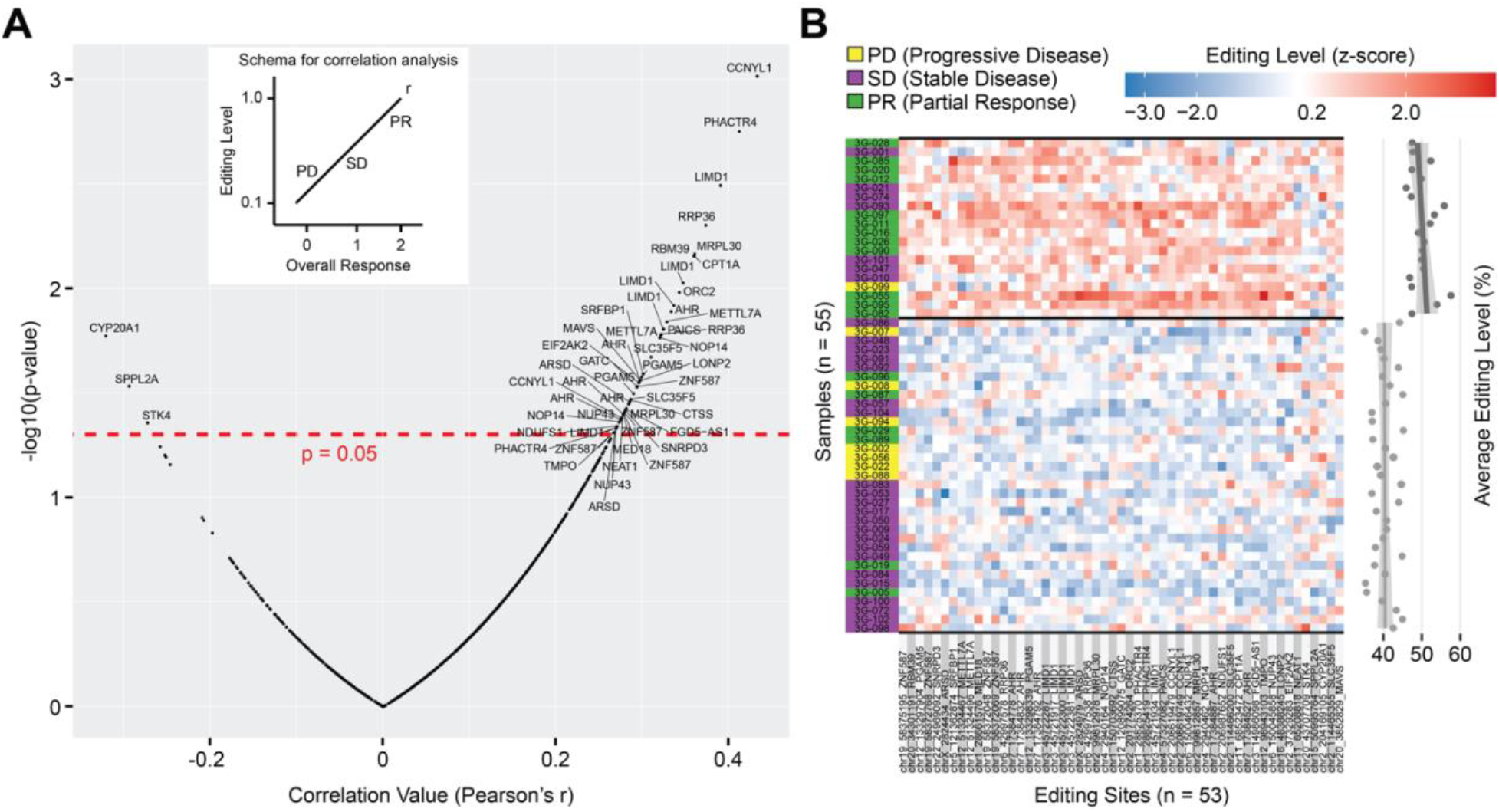
GCRE signature as a predictive marker in advanced GC. (*A*) Correlation of hotspot editing sites with overall response to chemotherapy. Categorical values of overall response are converted to numerical values (PD = 0, SD = 1, PR = 2), and then Pearson correlation test is applied to each site between editing levels and numerical values of overall response. A representative example of a positive correlation is illustrated by the inset plot. Of the cohort, 55 samples with tumour response data available are included in the analysis, n(PD) = 8, n(SD) = 29, n(PR) = 18. Gene symbols associated with the editing sites are shown for the significant cases (n = 53, p < 0.05), where genes with multiple sites are repeated. (*B*) k-means clustering of samples based on the RNA editing levels of 53 sites that significantly correlated with overall response (k = 2). Prediction accuracy of responders is 41/55 = 75%.

### GCRE score predicts responders with high accuracy

To predict chemotherapy outcome in a robust way, we derived a score based on the editing levels of 50 sites (GCRE signature) having positive correlation with the overall response (**Materials and Methods**). Briefly, we obtained the average z-score per sample based on GCRE signature (GCRE score), then we stratified the patients into 3 response groups (PD, SD, and PR) and predicted the responders based on this GCRE score. Overall, we observed a good performance (AUC = 0.77, **Figure 4A**), and at the cut-off value of 0.4, we achieved an accuracy of 84% (sensitivity = 67%, specificity = 92%) to predict the responders (**Figure 4B**), which was a better prediction compared to the clustering method (75%, **Figure 3B**). We also confirmed that reducing the number of editing sites or including 3 negatively correlated sites compromised accuracy (**Supplementary Figure 3**). Overall, these results suggest that employing the GCRE score based on the 50 sites contributing to the GCRE signature could stratify responders and non-responders to chemotherapy with high accuracy.

**Figure 4.**
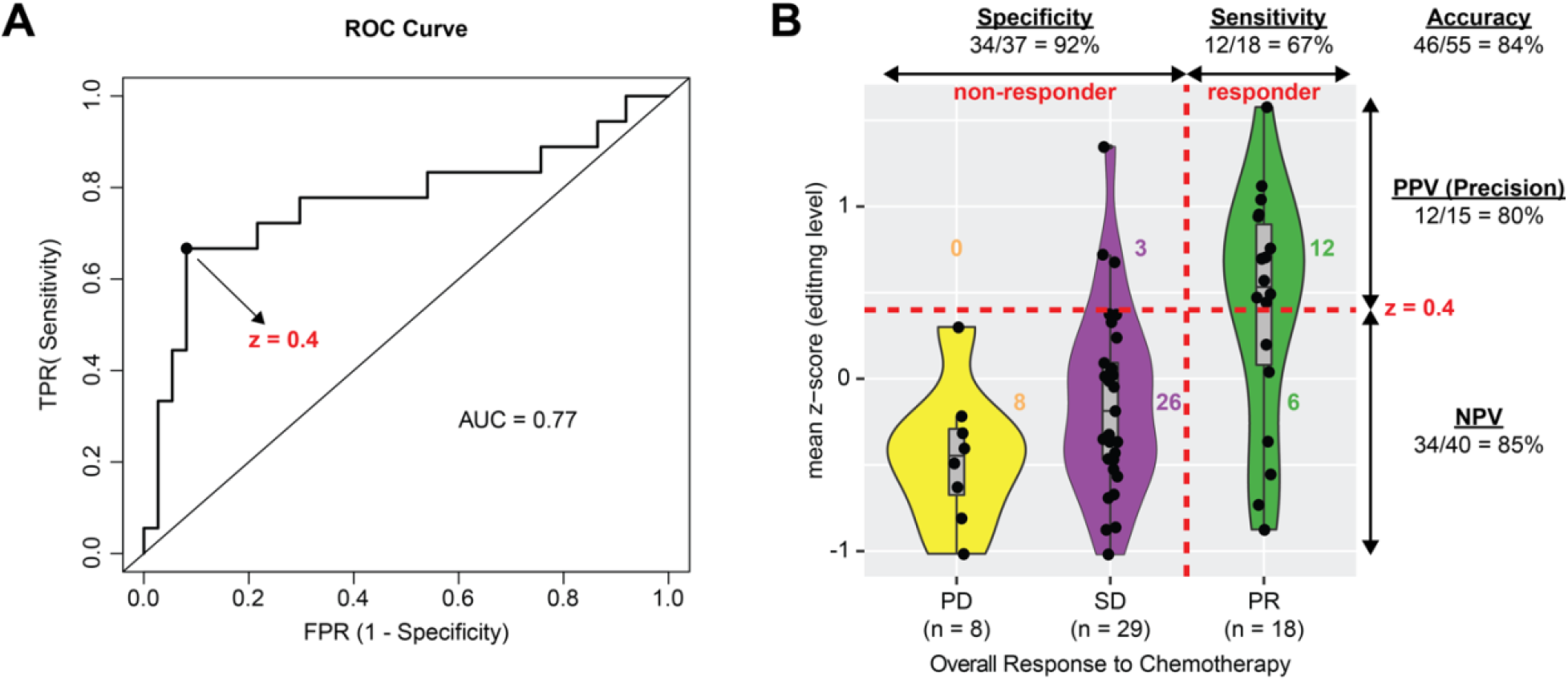
Utilisation of a GCRE score to predict chemotherapy response. (*A*) Receiver operating characteristic (ROC) curve showing the performance of GCRE score in prediction of responders. TPR = true positive rate, FPR = false positive rate, AUC = area under the curve. (*B*) Stratification of GC patients into chemotherapy response groups and prediction of responders based on GCRE score at the cut-off value of 0.4. GCRE score for each patient denotes average z-score of RNA editing levels across the panel of 50 sites in the GCRE signature. The statistical measures refer to the classification of responders and non-responders. PPV = positive predictive value, NPV = negative predictive value, non-responder = progressive disease + (PD) stable disease (SD), responder = partial response (PR).

### Validation of GCRE signature in GC cell lines

We next validated the GCRE signature in 6 commercially available GC cell lines. First, we assessed the drug response of each cell line to oxaliplatin by IC_50_ (the half maximal inhibitory concentration) values (**Figure 5A**) and further confirmed by foci formation assay (**Figure 5B**). Then, we quantified the RNA editing levels of 26 randomly selected sites from 50 sites of the GCRE signature in the same 6 cell lines by Sanger sequencing (**Figure 5C, Materials and Methods**). We found a negative correlation between the IC_50_ values and RNA editing levels (r = - 0.5), implying that GC cells with higher editing levels of the GCRE signature sites demonstrate higher chemosensitivity, which was consistent with our observation in the patient samples.

**Figure 5.**
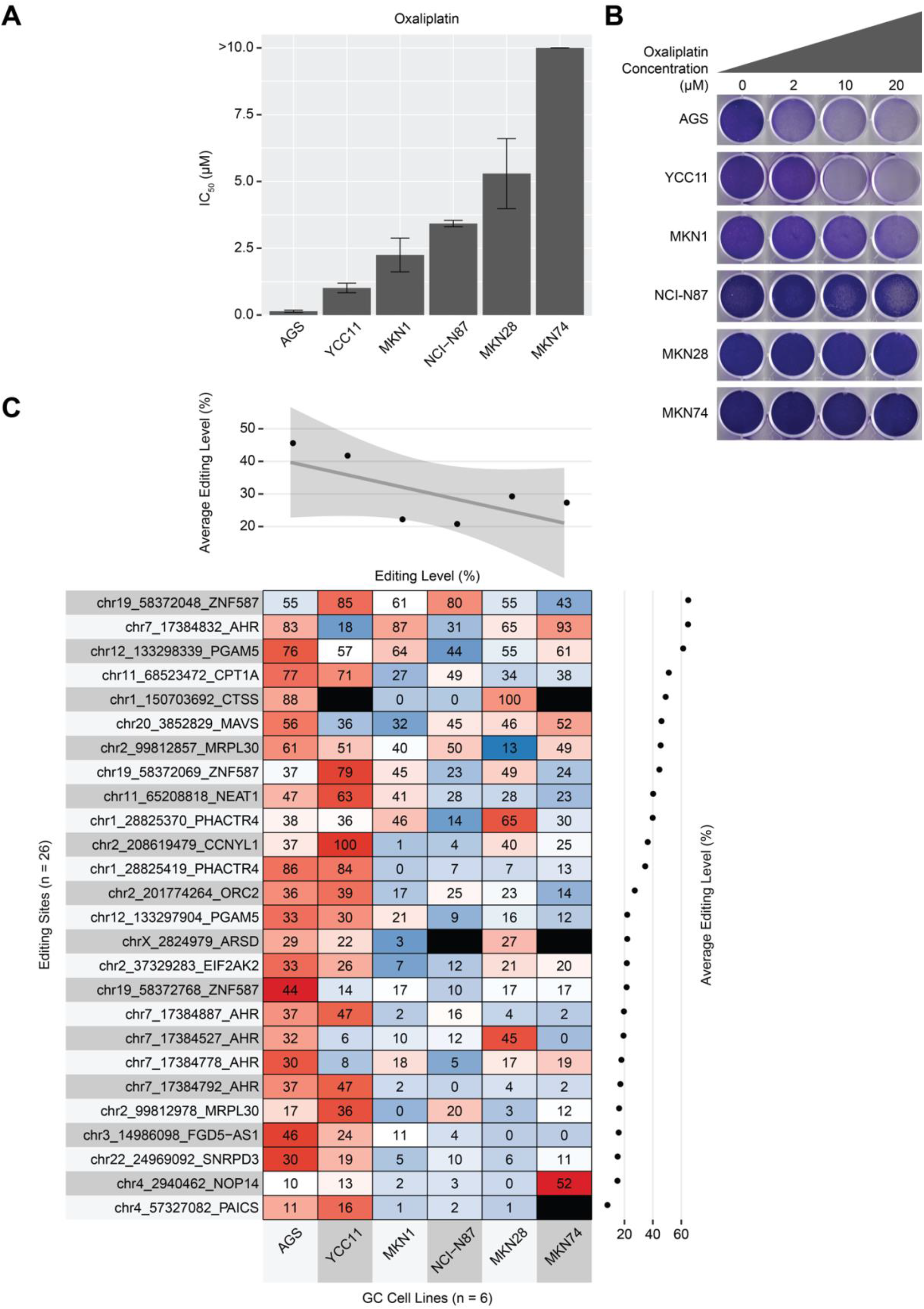
Validation of GCRE signature in GC cell lines. (*A*) Chemosensitivity of 6 GC cell lines to oxaliplatin. The bars show the average IC_50_ value of 3 biological replicates with standard error of means. (*B*) Foci formation assay of GC cell lines cultured with indicated concentration of oxaliplatin for 48 hours. Cells are stained with crystal violet. (*C*) Editing levels of 26 sites from the panel of GCRE signature quantified by Sanger sequencing. Numbers in cells denote the percentage of editing levels and colouring shows the relative gradient across the row (scale). Black cells denote undetectable editing level.

### GCRE signature is a prevalent in late-stage GC, independent of chemotherapy regimen

To infer whether our GCRE signature is representative of different tumor stages and chemotherapy regimens, we applied the GCRE score in TCGA STAD (stomach adenocarcinoma) cohort^6^. This cohort comprises of patients diagnosed with different stages of GC and treated with multiple combinations of drugs, along with multiple data points reporting for primary and follow-up treatment outcome (**Supplementary Table S5**). When we applied the GCRE score on these patients, we observed that responders in the latest stage (IV), but not in the previous stages (II and III), had significantly higher levels of editing in the panel of 50 sites compared to non-responders (**Figure 6**), confirming our observation in our advanced GC cohort, despite the diverse drug regimens between the two cohorts. We have excluded the patients at early stage (I) from the analysis as these patients most likely had undergone tumor removal rather than chemotherapy, so the response information was ambiguous. With available data, however, we could not validate the prognostic value of the GCRE signature possibly due to the small sample size (**Supplementary Figure 4**). These results suggest that the GCRE signature is coherent to very late stage of the disease as a predictive marker, independently of different chemotherapy regimens.

**Figure 6.**
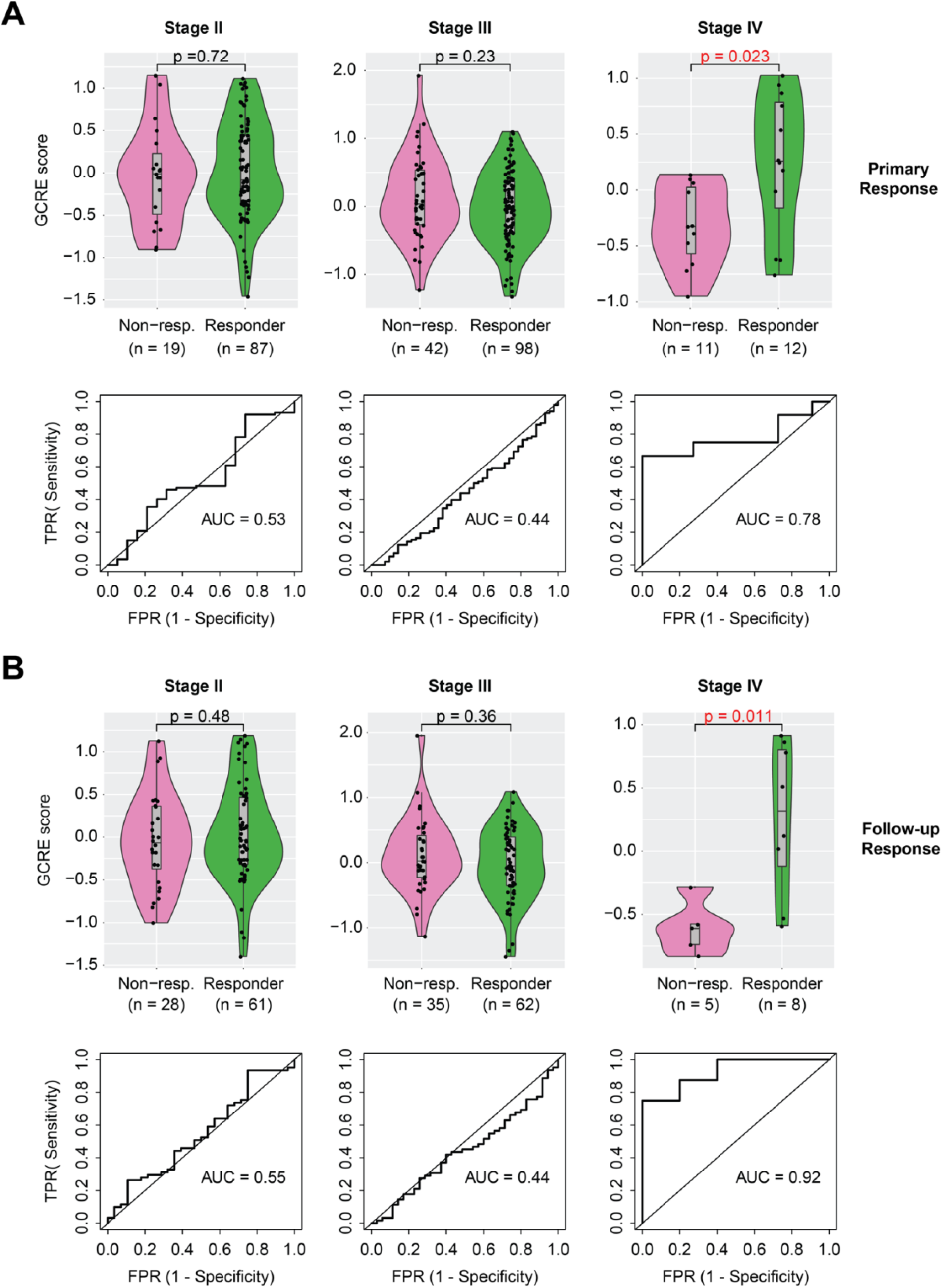
Validation of GCRE signature in TCGA STAD. Distribution (violin plots) and performance (ROC curves) of GCRE scores in TCGA STAD cohort stratified by disease stage and response type. For score calculation, 50 sites in GCRE signature and their editing levels in TCGA patients are used wherever available and at least 8 sites were required to be edited to calculate the score. Primary response (*A*) refers to the “primary_therapy_outcome” and follow-up response (*B*) denotes the “followup_treatment_success” as reported by TCGA. Response groups are merged based on the consensus of the individual treatment outcomes: Non-responder = stable/progressive disease, Responder = partial/complete remission, STAD = stomach adenocarcinoma, TPR = true positive rate, FPR = false positive rate, AUC = area under the curve. p-values are shown for Wilcoxon rank-sum test.

## Discussion

Patient stratification based on molecular information promises a great value for guiding diagnosis and treatment choice, particularly in genetically heterogeneous diseases such as gastric cancer. Till date, a number of studies attempted for the molecular classification of patients for this purpose, most of which are based on gene expression profiling utilizing microarray data^3–5, 7–9^. With the broader availability and better quality of next-generation sequencing data, now it became possible to reliably study other molecular features than gene expression. In this study, we propose A-to-I RNA editing as a novel molecular classifier in advanced GC, and show that an RNA editing signature can be used to stratify patients with differential benefit from chemotherapy, which may not be limited to GC and can be potentially extended to other cancer types and cohorts.

In recent studies, systematic and unbiased analysis of RNA editing led to the establishment of the driver role of individual editing events in several cancers^11, 12, 23, 24, 26, 27^. It also became evident that editing level of a considerable number of RNA editing sites is correlated with patient survival, indicating their utility as prognostic markers^11, 28, 38, 42, 43^. Several studies have also reported that a specific RNA editing event could selectively affect the outcome of cancer therapies. For example, protein-recoding RNA editing of *COG3* and *GRIA2* gene increases the drug sensitivity to MEK inhibitors^38^. In this study, we followed a top-down approach, starting from transcriptome-wide global editing events towards a small panel of sites with potential clinical utility. Accurate identification of RNA editing events rely on several factors, such as high sequencing quality, sufficient sequencing depth, rigorous filtering and strict inclusion criteria. In our dataset, we achieved a median of 117M reads per sample after adapter and quality trimming, so that we could apply a very high coverage threshold (>= 20 reads) to identify the editing sites. This led us to accurately quantify the differential editing levels across the samples, which was crucial for the high resolution of the clustering and correlation analyses.

Our approach demonstrates a simple yet powerful way of identification of key editing events in advanced GC, uncovering the GCRE signature consisting of 50 editing sites associated with 29 genes. Once clinically validated, this RNA editing signature can be conveniently managed in the laboratory setting for individual patients to assist in therapeutic decision-making.

In addition to the GCRE signature, we report a list of 780 editing sites shared by 104 advanced GC tumors, which presents a valuable resource for follow up studies. These sites include previously reported hotspots (frequently edited) with pathogenic role (e.g. *EIF2AK2, MAVS, GATC, CTSS, METTL7A*)^34, 42, 44–46^ as well as novel sites reported here for the first time (e.g. *NEAT1, ORC2, FGD5-AS1*). Although in this study we mainly focused on a small panel of 53 sites having significant correlation with the tumor response, the remaining sites potentially carry important information for GC pathology awaiting further exploration.

The availability of TCGA data gave us a great opportunity to validate the GCRE signature independently. The heterogeneity of these patients additionally allowed us to assess its specificity in terms of disease stage and treatment regimen, although the information on the type of chemotherapy for stage IV was limited. However, additional GC datasets comprising of diverse patient characteristics with available RNA-Seq and tumor response data are needed to conclude whether this signature is cohort-specific or representative of a certain patient class, and whether these 50 editing sites are definitive. In particular, the prognostic value of the GCRE signature needs investigation in cohorts with larger sample size.

Overall, in this study we have investigated the translational potential of RNA editing process in advanced GC by using transcriptomic and clinical data of a large cohort of patients. We discovered an RNA editing signature consisting of a small panel of genes independently of expression profiling, developed an RNA editing score which can predict responders with high accuracy, and validated them in GC cell lines and an independent cohort of TCGA. These findings suggest a novel clinical utility of RNA editing events for guiding chemotherapy treatment in this deadly cancer.

## Materials and Methods

### Gastric cancer cohort

A total of 104 patients from the 3G Trial^7^ were involved in this study. The patients were diagnosed with metastatic or recurrent GC and enrolled onto the first line palliative chemotherapy (platinum-fluoropyrimidine doublet chemotherapy regime). For each eligible patient, fresh endoscopic biopsies of the primary tumor *in situ* within 3 weeks prior to treatment initiation were used for RNA-Seq analysis. Treatment response to chemotherapy (PD: progressive disease; SD: stable disease; PR: partial response) was assessed by radiologists who were blinded to the study. At the cut-off date, in this cohort of 104 patients, overall survival and tumor response data had been obtained for 54 and 55 patients, respectively, where a total of 50 patients had both OS and tumor response data.

Samples were collected from the cancer centres in Singapore (n = 37) and South Korea (n = 67) between 2010 - 2018. A written informed consent was obtained from all the patients prior to the enrolment to the trial, and the study was done in accordance with the Declaration of Helsinki and International Conference on Harmonisation and Good Clinical Practice guidelines. The protocol was approved by the Institutional Review Board at each study site and complied with local laws and regulations.

### RNA sequencing

A total of 1 μg of total RNA was used to create libraries with Illumina TruSeq Stranded Total RNA Library Prep Kit (Illumina) according to manufacturer’s instructions. Library fragment size was determined using the DNA 1000 Kit on the Agilent Bioanalyzer (Agilent Technologies). Libraries were quantified by qPCR using the KAPA Library Quantification Kit (KAPA Biosystems). Libraries were pooled in equimolar and cluster generation was performed on the Illumina cBOT system (Illumina). Sequencing (150bp pair-end) was performed on the Illumina HiSeq 3000 system at the Duke-NUS Genome Biology Facility, according to manufacturer’s protocol (Illumina).

### Identification of RNA editing events

A bioinformatics pipeline adapted from a previously published method^47^ was used to identify RNA editing events from RNA-Seq data by using the CSI NGS Portal^48^ (https://csibioinfo.nus.edu.sg/csingsportal). For each sample, raw reads with adapters were trimmed by using Trimmomatic (v0.38)^49^ retaining the reads with >= 35 bases and average read quality score > 20 after trimming. Clean reads were mapped to the reference human genome (*hg19*) with a splicing junction database generated from transcript annotations derived from UCSC^50^, RefSeq^51^, Ensembl^52^ and GENCODE (v19)^53^ by using Burrows-Wheeler Aligner with default parameters (*bwa mem*, v0.7.17-r1188)^54^. To retain high quality data, PCR duplicates were removed (*samtools markdup -r*, v1.9)^55^ and the reads with mapping quality score < 20 were discarded. Junction-mapped reads were then converted back to the genomic-based coordinates. An in-house perl script was utilized to call the variants from samtools pileup data and the sites with at least two supporting reads were initially retained. The candidate events were filtered by removing the single nucleotide polymorphisms reported in different cohorts (1000 Genomes Project^56^), NHLBI GO Exome Sequencing Project (https://evs.gs.washington.edu/EVS/), dbSNP (v150)^57^) and excluding the sites within the first six bases of the reads caused by imperfect priming of random hexamer during cDNA synthesis. For the sites not located in Alu elements, the candidates within the four bases of a splice junction on the intronic side, and those residing in the homopolymeric regions and in the simple repeats were all removed. Candidate variants located in the reads that map to the non-unique regions of the genome by using BLAST-like alignment tool^58^ were also excluded. At last, only A-to-G editing sites based on the strand information from the strand-specific RNA-Seq data were considered for all the downstream analyses. The genomic regions of the editing variants and the associated genes were annotated by using ANNOVAR (v2018)^59^ with the UCSC *refGene* table annotation^60^. We applied the same pipeline on TCGA STAD (The Cancer Genome Atlas - stomach adenocarcinoma) cohort^6^.

To identify high confidence and common editing events, stringent filtering criteria were applied. Specifically, each editing site was required to have a coverage of at least 20 reads and editing frequency higher than 0.1 (10%) in all the samples. This resulted in 780 high confidence editing sites shared by 104 samples of our GC cohort (**Figure 1C**). For TCGA STAD cohort, we did not apply these thresholds for the validation of GCRE signature in order to include more sites, as the number of high-confidence editing sites were relatively fewer in TCGA due to the lower sequencing depth. In TCGA, we included only those samples with at least 8 out 50 sites in the GCRE signature were found to be edited (**Figure 6**).

### Clustering analyses and heatmaps

The clustering and heatmap analyses were performed by using R package “superheat”^61^. The RNA editing levels of the corresponding sites were used as the distance matrix to perform the k-means clustering for the heatmaps in an unsupervised manner (scale = TRUE, n.clusters.rows = 2, clustering.method = “kmeans”). The side plots were drawn using “scattersmooth” parameter with the “lm” method based on the average editing value of the corresponding rows or columns calculated by “rowMeans2” and “colMeans2” functions from the “matrixStats” package, respectively.

### Correlation of RNA editing with response to chemotherapy

Pearson correlation analysis was performed by using “cor.test” function in R with “pearson” method. For each of 780 editing sites, Pearson correlation coefficient (r) and associated p-value were calculated between RNA editing levels and the overall response to chemotherapy across 55 patients with available data. The editing frequencies calculated from the RNA-Seq data were used as the first vector. As the second vector, the overall response data was used after transforming the original categorical variables into numerical variables (PD = 0, SD = 1, PR = 2) so that the correlation analysis can be performed. P-values were used to assess the editing sites with significant correlations at p < 0.05 threshold, which resulted in 50 positively and 3 negatively correlated sites (**Figure 3A**).

### GCRE score calculation and GCRE signature

To predict the chemotherapy outcome based on RNA editing, we developed a GCRE score based on the “z-score” using 50 sites that showed significant positive correlation with the overall response. First, z-transformation was performed for each site based on the RNA editing levels. Then the samples were ranked by using the average z-score across the sites. The samples above and below a cut-off value (0.4) were regarded as the high and low editing groups, respectively. The statistical measures were calculated based on the prediction of the responders in the cohort.

### Gastric cancer cell lines and validation of the GCRE signature

A total of 6 cell lines were used for the validation of the GCRE signature. AGS and NCI-N87 were purchased from the American Type Culture Collection (Manassas, VA). MKN28, MKN1, and MKN74 were obtained from Japanese Collection of Research Bioresources Cell Bank. YCC11 cell line was provided by Singapore Gastric Cancer Consortium (SGCC). All the cell lines were cultured in Roswell Park Memorial Institute (RPMI) medium (Gibco BRL, Grand Island, NY, USA) supplemented with 10% fetal bovine serum (FBS) (Gibco BRL). Cells were grown in a humidified incubator with 5% CO_2_ at 37°C.

The editing levels of 26 out of 50 editing sites in these 6 cell lines were quantified by Sanger sequencing. The sites were selected as the top 10 sites which showed the highest change in editing level between high and low editing group as defined in the RNA-Seq data, and additional 16 sites that are randomly picked from the remaining sites in the panel of GCRE signature. Drug response of the cell lines to oxaliplatin (Sigma-Aldrich) were assessed by IC_50_ values using MTT (3-(4,5-Dimethylthiazol-2-yl)-2,5-diphenyltetrazolium bromide) assay. Briefly, GC cell lines were seeded in 96-well plates with 2.5 × 10^3^ to 10 × 10^3^ cells/well according to their growth rate. After 72-h oxaliplatin drug treatment, 10 μL MTT substrate (Sigma-Aldrich) was added into each well followed by 3-hour incubation. MTT substrate was then removed and cells were lysed by addition of 100 μL MTT stop solution. Absorbance at 570 nm was measured using Tecan microplate reader.

For the foci formation, cells were seeded in 6-well plate with 3 × 10^4^ to 15 × 10^4^ cells/well and cultured with indicated concentration of oxaliplatin for 48 hours. Cells were stained with crystal violet (Sigma-Aldrich) for colony visualization.

### Statistical analyses

As a rule of thumb, all the available features and the samples were included in the respective analyses as long as the data being investigated were non-missing. Initial clustering analysis was performed in an unbiased manner based on the editing levels of all the sites identified in all the samples. The downstream analyses were performed by using a panel of 50 sites that are selected solely based on the correlation value of their editing levels obtained from RNA-Seq data and the overall response to chemotherapy assessed by clinicians and radiologists. GCRE score was derived based on the editing levels of the selected genes in all the samples with response data available. No manual selection or exclusion was applied on the genes, editing sites or samples for this study, unless limited by the availability of data under investigation, e.g. 55 out of 104 samples had response data. These criteria were also applied to the cell lines in the same way as per patient samples. For the experimental validations, unless otherwise indicated, the data are presented as the mean ± standard error of mean (SEM) of three independent experiments. Wilcoxon rank-sum test was applied when comparing distributions, Pearson correlation coefficient was reported for paired correlation analyses, and log-rank test p-value was shown for the survival analyses. A p-value of less than 0.05 was considered to be statistically significant, and the type of the statistical test applied was indicated appropriately.

### Data availability

RNA-Editing pipeline is available online at the CSI NGS Portal^48^ (https://csibioinfo.nus.edu.sg/csingsportal). The bioinformatics code for the downstream analyses are available upon reasonable request. The processed data are available as **Supplementary Tables**.

## Author Contributions

Conception, design and supervision: C.L. (RNA editing analysis), J.B.-Y.S., W.P.Y. and S.Y.R. (3G trial)

Analysis and interpretation of clinical data: R.S. and M.C.H.N.

RNA-Sequencing of clinical samples: P.T., L.M.H., T.S.T., O.X.W., A.T.L.K. and E.T.

Bioinformatics analyses: O.A. conceived and performed all the bioinformatics analyses, with inputs from H.Y.

Cell-based experiments: Y.S. and X.Y.K. performed all the wet lab experiments.

Manuscript writing: O.A. and C.L. wrote the manuscript.

All authors read and approved the final version of the manuscript.

## Acknowledgements

We greatly thank to the patients and their families for participating in the clinical trial and for contributing tissue magnanimously to the biomarker study.

## Abbreviations

A-to-I: adenosine-to-inosine
AUC: area under the curve
GC: gastric cancer
IC_50_: the half maximal inhibitory concentration
PD: progressive disease
PR: partial response
RE: RNA editing
RNA-Seq: RNA sequencing
SD: stable disease
SGCC: Singapore Gastric Cancer Consortium
STAD: stomach adenocarcinoma
TCGA: The Cancer Genome Atlas.

## Supplementary Information

**Supplementary Tables** (Please refer to the Excel file)

**Table S1** - Clinical data of the GC cohort (3G Trial)

**Table S2** - The 780 shared editing sites across 104 patients (hotspots)

**Table S3** - Cox Proportional-Hazards for editing clusters and baseline patient characteristics

**Table S4** - The 53 GCRE signature sites

**Table S5** - Clinical data of TCGA STAD cohort

**Supplementary Figure 1.**
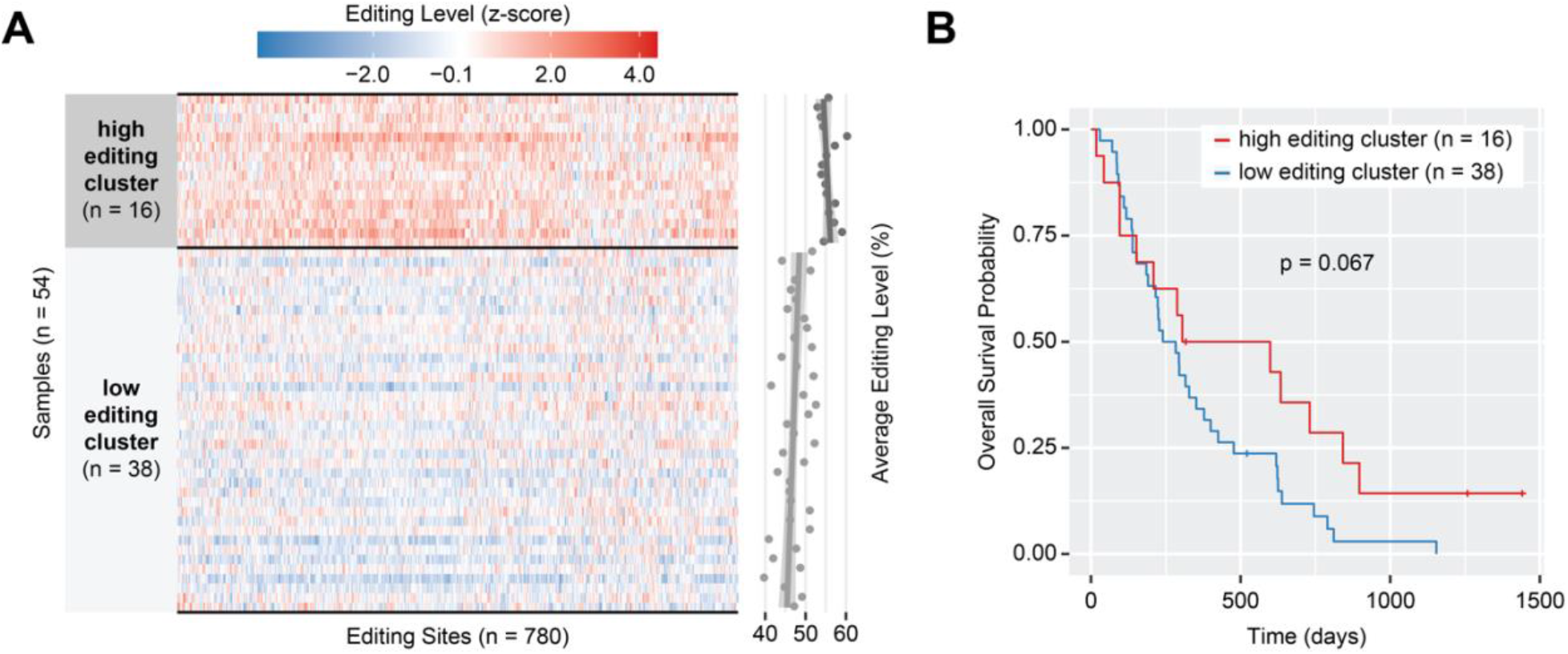
RNA editing hotspots as a prognostic marker in advanced GC. (A) Hierarchical clustering of GC samples based on RNA editing levels of 780 hotspot editing sites. The scatterplot shows average editing levels per sample. Of the cohort, 54 samples with survival data available are included in the analysis. (B) Survival plot of the two editing clusters.

**Supplementary Figure 2.**
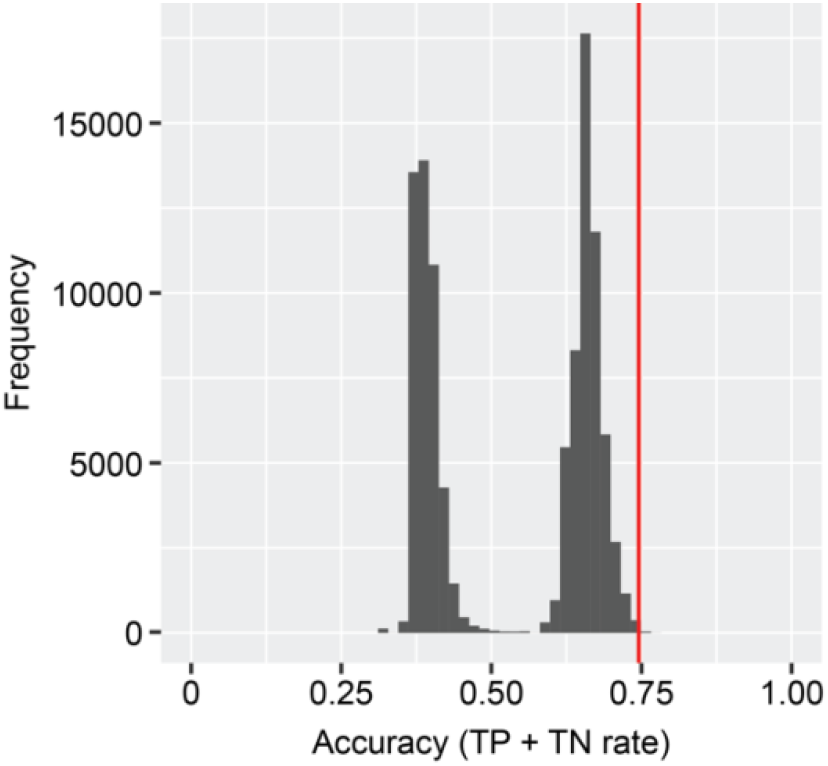
A randomization test of GCRE signature. Histogram of prediction accuracies for chemotherapy response in a randomized control. The test is performed by hierarchical clustering of 55 samples based on the editing levels of randomly picked 53 sites out of 780 hotspot editing sites repeated for 100,000 times. In each round, the test statistic is calculated as the prediction accuracy of responders in the clusters. The observed accuracy of the actual GCRE signature derived from the correlation test is highlighted with the red line, where empirical p-value = 0. 00409. TP = true positive, TN = true negative.

**Supplementary Figure 3.**
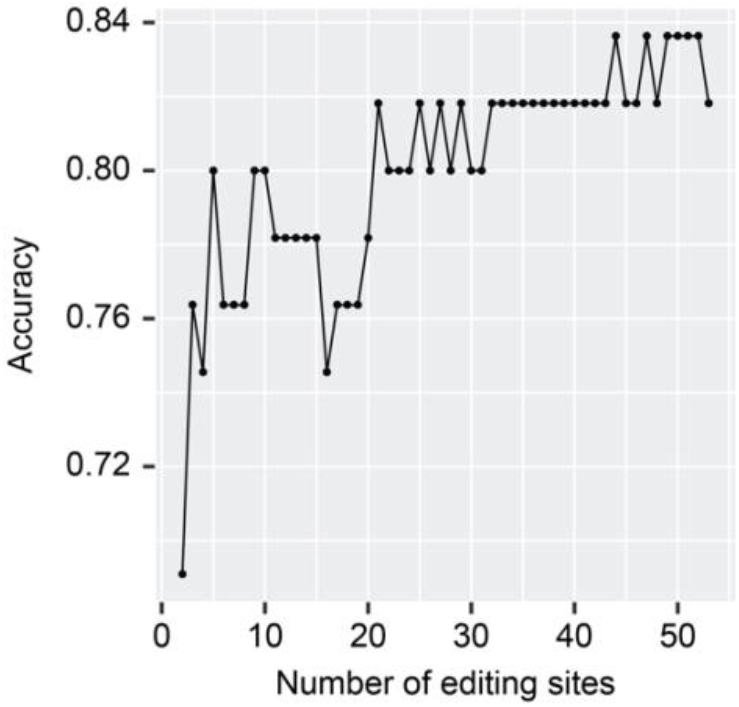
Selection of optimal number of editing sites for GCRE signature. Accuracy distribution based on cumulative number of editing sites. The sites are ranked based on the difference in editing level between responders and non-responders, and accuracy is calculated for increasing number of sites from 2 to 53. The first 50 sites are the positively correlated sites whereas the last 3 are the negatively correlated sites with the overall response, where accuracy is steadily highest around 50 sites.

**Supplementary Figure 4.**
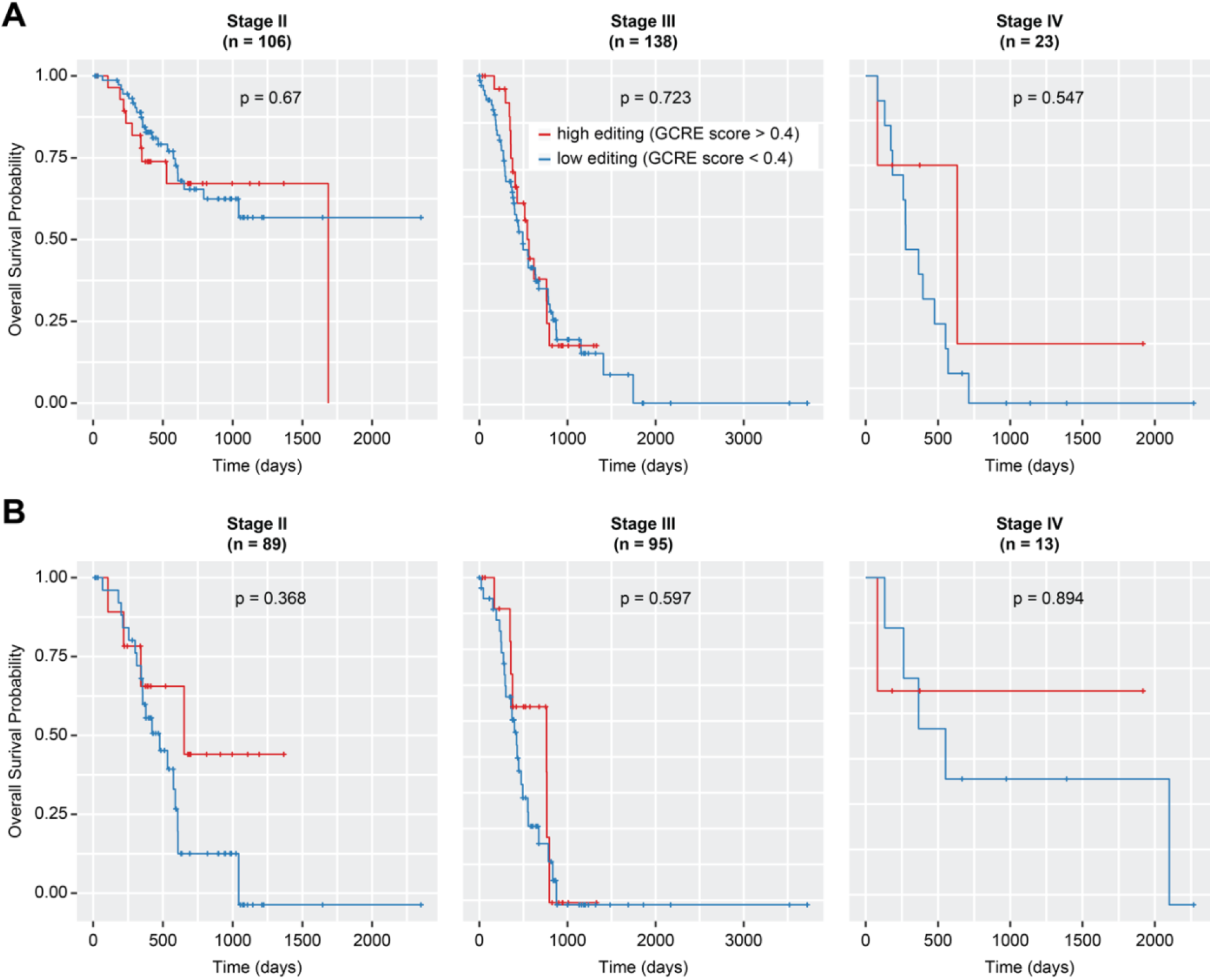
Survival plots of patients with high vs low editing in TCGA STAD cohort stratified by disease stage and response type. High and low editing was defined as the GCRE score higher and lower than 0.4, respectively. The total number of patients correspond to those in **Figure 6**, with 2 patients missing survival information in Stage III.

